# Implementation of Design of Experiments (DOE) for Optimization of Feeding Strategy and Glyco-Engineering of Trastuzumab Biosimilar

**DOI:** 10.1101/584144

**Authors:** Rasoul Mahboudi, Sepideh Samavat, Amir Afrah, Mehdi Khorshidtalab, Arezou Fadaei Tehran, Paria Motahari, Farnoush Jafari Iri Sofla, Shayan Maleknia

## Abstract

Fed-batch cell culture is the most commonly used process for antibody production in biopharmaceutical industries. Basal media, feed, feeding strategy and glycan structures are always among the most important concerns during process development and optimization. In this study, first, a traditional screening study was performed to identify the top media/feed combinations by evaluating the cell culture performance including cell growth and protein titre. Optimization of the process was also performed using response surface methodology in order to find the most optimum feeding strategy and glucose set point regarding final titre of the recombinant monoclonal antibody being produced in Chinese hamster ovary cell line. The focus of this study is not only on titre, but also on product quality and comparability especially protein glycosylation. The prediction model of product titre as a function of feeding percentage and glucose set point was successfully applied for the second set of experiments that was performed for glycan improvement. Statistical design of experiments was applied to determine the most important factors and their effects on galactosylated and afucosylated glycans. Uridine, manganese, galactose and fucosyltransferase inhibitor were chosen to evaluate if their presence can affect glycans and to obtain their best combination for fed-batch culture supplementation. We determined that 2.5 % daily feeding combined with maintaining the glucose set point on 2.5±0.2 g/L could achieve final titre of 2.5± 0.1 g/L. Galactosylation of antibody was increased about 25% using MnCl_2_ and galactose while afucosylation was increased about 8% in presence of fucosyltransferase inhibitor. Galactose and Mn^2+^ led to a shift from G0F to G1F and presence of Fucosyltransferase inhibitor caused to an increase in G0 compared to its absence. These results demonstrated that supplementation of culture with all these components can provide exact control of antibody galactosylation and fucosylation with minimal impact on culture characteristics and product quality attributes. Subsequently, validation experiments were also carried out in 5L STR bioreactors which showed that similar results could be achieved in bioreactors compared to shake flasks regarding both titre and quality.

## Introduction

Monoclonal antibodies (mAbs) have become famous therapeutic agents in treatment of cancer, inflammatory, respiratory and infectious diseases. For mAb based therapies, typically, high doses in the range of hundreds of milligrams to a gram are needed per dose for achieving the favored therapeutic effect. In addition, high doses of antibodies are required over a long period of time, which requires large amounts of purified protein. Manufacturing capacity and cell line productivity are among most important factors that can affect the cost and efficacy of drug substance production (1). Therefore, increasing the productivity of antibody-producing cells in biopharmaceutical industries to meet the market demand is a real challenge (2).

Most monoclonal antibodies production platforms are based on firmly established fed-batch culture mode due to its ease of scale up and compatibility with large scale manufacturing of up to thousands of litres. The volumetric productivity of such cultures was typically improved from 50 mg/L/d in 1990s to about more than 200 mg/L/d at present (3). Basal and feed media are two components of fed-batch process that caused to cell growth and production, respectively by providing essential nutrients during the batch. For fed-batch mode, a sequential approach is conventionally used in which an optimal basal media is first chosen according to the growth characteristics of the cell line and quality attributes of the protein being produced. In next step a feed medium is also optimized regarding the type of feed and the feeding strategy in a way to provide the highest titre while maintaining the critical quality attributes in their acceptable range (4).

Traditional and statistical methods can be used for efficient selection of cell culture media/feeds combination and feeding strategy optimization. Traditional methods are costly and time consuming and do not seem favorable for biopharmaceutical companies which are looking forward to reduce time to market. Nowadays, approaches that combine high-throughput screening platforms with statistical design of experiment (DOE) such as factorial design and response surface methodology (RSM) are commonly used in biopharmaceutical companies to reduce the bioprocess development time and cost in Research and Development units (5). Factorial designs are used in order to find impact of several factors on a single response. However, RSM allows to achieve the most optimum response with selected factors.

Glycosylation is one of the most important post-translational modifications and critical quality attributes of therapeutic glycoproteins including monoclonal antibodies. These glycan chains on Fc region of mAbs are essential for their efficacy, stability and immune effector functions i.e. antibody-dependent cell-mediated cytotoxicity (ADCC) and complement-dependent cytotoxicity (CDC) (6). In particular, afucosylation and galactosylation play critical roles in ADCC and CDC, respectively (7). Additionally, during the development of a biosimilar drug product, it is essential to completely demonstrate similarity in both therapeutics regarding their critical quality attributes (CQAs). Similarity of the quality attributes between both innovator and biosimilar products should be proven before the new drugs entered to the market (8). Monoclonal antibody glycosylation can be modulated using various commercial products that were designed to enhance glycosylation. Among these additives, EX-CELL Glycosylation Adjust supplement (Gal+) (MilliporeSigma) has been shown effectivee3 results regarding enhanced glycosylation (9).

In this study, according to the interrelated behavior of feed and media on cell culture process parameters, combined optimization of basal and feed media was performed in order to develop a process that results in highest productivity of a CHO cell line producing an anti-HER2 IgG1 monoclonal antibody (trastuzumab biosimilar). After finding the best basal/feed combination through conventional media/feed selection strategies, RSM methodology was used for finding the optimum level of feed and glucose set point for achieving the highest titre of the monoclonal antibody. Another important aim of this study was fine tuning of the culture composition through supplementation of nucleotide sugar precursors and glycosyltransferase enzymes cofactors or inhibitors to optimize galactosylated and afucosylated content of the antibody being produced. For increasing antibodies with decreased amount of fucosylation i.e. afucosylation, a fucosyltransferase inhibitor 2F-peracetyl-fucose was used (10). In order to increase the level of galactosylated forms, three factors that were expected to potentially enhance intracellular galactosylation, namely galactose, uridine and Mn^2+^ were supplemented to the culture. Galactose and uridine are the main components for production of UDP-gal (a nucleotide sugar precursor that is added to the oligosaccharide moieties being present on Fc region of antibodies) while Mn^2+^ is the cofactor of galactosyltransferase enzyme that adds galactose residue to the glycan chains (11). The optimum concentration of all these supplements added to culture for improving glycosylation were fine-tuned by performing experiments that were designed through Full Factorial design of experiments. Finally, the models were further verified by performing a series of experiments in pre-determined optimum conditions predicted by the models.

## Materials and methods

### Basal media and feed selection

The CHO cell line producing trastuzumab biosimilar was kindly provided by AryoGen Pharmed Inc. (Alborz, Iran). This cell line was derived from CHO-S cells (Gibco, Catalog No. A11364). In order to find the best combination of basal media and feed that could result in optimum cell growth, and protein productivity, fed-batch cultures were performed by using commercially available basal/feed media combinations. Cell growth characteristics and protein titre were compared in six fed-batch culture conditions with different basal/feed combinations. Three chemically defined basal media were compared being: 1. CD1, 2. CD2 and 3. CD3 media, with media 1, 2 purchased from (Lonza, Verviers, Belgium) and medium 3 purchased from UGA Biopharma GmbH (UGA, Hennigsdorf, Germany). Two chemically defined feed systems were used in combination with three basal media mentioned above and compared. The feed systems used include: 1. FC from Life Technologies; and 2. FA and FB from GE Healthcare. Glucose concentrations of FC and FA was about 30 and 80 g/L, respectively. Media and feed solutions and supplements were all prepared according to manufacturer’s protocols. All cultures contain 0.1% (v/v) Anti-clumping agent (Lonza, Verviers, Belgium).

One vial of research working cell bank was thawed in 500 ml shake flask containing 100 ml pre-warmed media. Cells were sub-cultured every 3 days to initial cell density of 0.6 ±0.05×10^6^ cells/ml for at least 6 passages. Then fed-batch cultures were started with initial working volume of 100 ml in 500 ml shake flasks with the same initial cell density. Cells were incubated in an incubator with 5% CO_2_ at 37 °C on orbital shaker with 100 rpm shaking speed and 30 mm orbital shaking diameter. In all culture conditions feeding was started on 2^nd^ day of culture with a daily bolus addition according to Table 1. Batches were harvested on either day 15 or when viability dropped below 70%, which one occurred earlier.

**Table 1.** Feeding protocols used in basal/feed selection experiments using three different basal media and two feeds.

| Basal media | Start feed (day) | Feed 1 (day, V/V %) | Feed 2 (day, V/V %) |
| --- | --- | --- | --- |
| CD1 | 2 | FA (2%), daily | FB (0.2%), daily |
| CD1 | 2 | FC (2%), daily | — |
| CD2 | 2 | FA (2%), daily | FB (0.2%), daily |
| CD2 | 2 | FC (2%), daily | — |
| CD3 | 2 | FA (2%), daily | FB (0.2%), daily |
| CD3 | 2 | FC (2%), daily | — |

### Integrated medium and feeding strategy optimization

In a typical fed-batch culture, the basal medium usually supports cell growth, while the feed medium contributes in increasing the productivity. After determining the top basal media/feed combination, the next development phase was started to improve the feeding strategy in order to fine tune the batch duration combined with final expression. To optimize the feeding strategy, two independent variables of feed percent (factor A) and glucose set point (factor B) were chosen to see their effect on protein titre (response). The effect of these two parameters on titre response was analyzed by standard response surface methodology (RSM) design called central composite design (CCD). RSM statistical method is suitable for optimization of the process parameters with a minimum number of experiments, as well as finding the quadratic and interaction effects between factors. Minitab version 17 (Minitab, Inc.) was used to design the experiments. Each factor was coded as −1 (low level), 0 (center point) and +1 (high level) and their uncoded values are represented in Table 2. These two factors were chosen since their influence on culture characteristics were known according to our previous knowledge. Central composite design with a total of 13 experiments was used to obtain a quadratic model. The experimental run was randomized in order to omit the error and effect of the uncontrolled factors. The experiments being performed with center point values were repeated five times in order to have a good estimation of the experimental error. The model terms with *p-value* > 0.05 (if present) were removed in order to obtain a model with more adequacy.

**Table 2.** The amount of different factors chosen for experiments in Central Composite design.

| Factors | Low level (-1) | Centre point (0) | High level (+1) |
| --- | --- | --- | --- |
| Feed percent (%) | 2 | 2.5 | 3 |
| Glucose set point (g/L) | 2 | 3 | 4 |

**Table 3.** Characteristics of different fed-batch cultures regarding basal/feed media combinations.

| Basal-media combinations/<br>characteristics of batches | CD1 |  | CD3 |  | CD2 |  |
| --- | --- | --- | --- | --- | --- | --- |
|  | FA, FB | FC | FA, FB | FC | FA, FB | FC |
| Batch duration (day) | 15 | 14 | 15 | 15 | 15 | 13 |
| Max cell density ($\times 10^6$ cells/ml) | $14.4 \pm 0.4$ | $11.2 \pm 0.2$ | $16.3 \pm 0.4$ | $13.3 \pm 0.2$ | $12.1 \pm 0.3$ | $8.7 \pm 0.2$ |
| Final titre (mg/L) | $580 \pm 26$ | $394 \pm 30$ | $1426 \pm 30$ | $743 \pm 40$ | $316 \pm 20$ | $173 \pm 15$ |
| Cell specific productivity (Qp)<br>(pg/cell/day) | $4.3 \pm 0.1$ | $4.0 \pm 0.0$ | $8.9 \pm 0.0$ | $6.1 \pm 0.1$ | $3.0 \pm 0.1$ | $2.2 \pm 0.0$ |

### Glycan engineering experimental designs

Experimental data was generated according to a 2^4^ factorial design. Full factorial design was chosen in order to include all possible combinations of factors at selected levels. The factors and their high and low limit ranges were MnCl_2_ (Sigma, life science) (0-40 µM), galactose (Applichem, USA) (0-50 mM), Uridine (Sigma, life science) (0-4 mM), fucosyltransferase inhibitor 2F-peracetyl-Fucose (merck, calbiochem) (0-30 µM). Uridine was supplemented on days 6 and 8 of the culture while, MnCl_2_, galactose and fucosyltransferase inhibitor (FTI) were added to basal media prior to inoculation of cells. CHO cells were cultured in 500 ml shake flasks with an effective volume of 100 ml, incubated at 37 °C with a 5% CO_2_, and agitated at 80 rpm. Each shake flask was inoculated with an approximate cell density of 0.6 ± 0.05×10^6^ cells/ml. All experiments were performed with CD3 media and FA, FB feeds. Feeding strategy was set on 2.5% FA and 0.25% FB starting from day 2 till day 14 of the batch. Furthermore, glucose (Merck, Germany) was supplemented to keep on 2.5 ±0.2 g/L using 400 g/L glucose stock solution. Cell density, viability, pH and lactate were measured daily. Protein titre was also measured on days 3, 5, 7, 9, 11, 13, and 15 of the batches using Protein A HPLC method. All the conditions were harvested on 15^th^ day of the batch and were purified using MabSelect SuRe LX resin. Glycan analysis was performed and the data set was used for analyzing the factorial design in Minitab software. Three responses considered for this experimental design included: final titre (mg/L), Sum of oligosaccharides with galactose and without fucose. Total of 17 runs were designed using general full factorial design considering one center point for each factor being tested. Factor coefficients and *p-values* were determined after analyzing the design using ANOVA.

### Cell density and viability determinations

Sampling of cultures were performed daily by taking 1 ml sample from each shake flask. Cell density and cell viability were determined by Trypan blue exclusion assay using a Neubauer cytometer. Glucose and lactate concentrations were measured daily with a BioProfile Analyzer 400 (Nova Biomedical, Barcelona, Spain).

### Monoclonal antibody quantification

The concentration of the monoclonal antibody in cell culture supernatant samples was determined by the Mab Pac protein A affinity column (Thermo scientific, CA, USA). Two buffer solutions were used for HPLC test. The equilibration buffer or mobile phase A (50 mM PBS, 150 mM sodium chloride, (Merck, Germany) and 5% acetonitrile (Merck, Germany) pH 7.5. The mobile phase B was the same as mobile phase A buffer, with the only difference that pH of this solution was adjusted on 2.5 by Orthophosphoric acid. The elution gradient was programmed from 0% to 100% of elution buffer B in approximately 5 min. The standard curve generated with the purified monoclonal IgG with predefined concentrations and the mAb quantification was performed based on this curve.

### Antibody purification

Product quality assays were performed on samples purified by Protein A chromatography using MabSelect SuRe LX resin (GE Healthcare, Amersham, UK). The column was equilibrated in a citrate/sodium chloride buffer at pH 7. Cold, clarified bioreactor supernatant samples were loaded directly onto the column, washed with citrate/sodium chloride buffer at pH 7, eluted in citrate buffer at pH 3.5, and neutralized using a Tris buffer.

### Glycan analysis

N-linked glycans were released from purified antibody samples by overnight incubation with N-Glyccosidase F (Prozyme, GKE-5006B). The proteins were removed using LudgerClean glycan EB-10 cartridges (Ludger, Oxfordshire, UK). The oligosaccharides were labeled with the fluorophore 2-aminobenzamide (2AB) using Ludger 2-AB (2-Aminobenzamide) glycan labeling kit (Cat No. LT-KAB-A2). The excess 2AB fluorophore was removed using LudgerClean S glycan cleanup cartridges. 2-AB Labeled glycans were eluted from the cartridges using ultra purified water (UPW) and were incubated in 5 °C for 24 h. The samples were then resuspended using 80% Acetonitrile solution and a sample was injected in to ACQUITY UPLC Glycan BEH Amide 1.7 µm (2.1×150 mm) Column equilibrated in Acetonitrile and eluted with a gradient of 100 mM Ammonium formate pH 4.5 ± 0.1. The two mobile phases used were solvent A (0.1 M Ammonium formate pH 4.5 ± 0.1) and solvent B (Acetonitrile). Elution of the sample was performed while Fluorescence was set at Ex = 330 nm and Em = 420 nm. The peaks that are present in the UPLC chromatogram of Herceptin biosimilar as well as innovator drug are as follows: G0FGN, G0, G0F, Man5, G1, G1F, G1F’ and G2F. The amount of each structure is expressed as the percentage of total peak area. The amount of galactosylated structures is the sum of G1, G1F, G1F’ and G2F peaks, while the afucosylated population is the sum of G0 and G1 peaks area. Man5 structure is the only structure that represents the amount of oligosaccharides with mannose.

### Validation and verification of the models

Both experimental designs were verified by performing 5 runs in 5-L STR bioreactors in order to demonstrate the validity and predictive ability of both models obtained in this study. The culture process in bioreactors started by inoculation of 0.6 ±0.05 ×10^6^ cells/ml using the same media/feed used in small scale experiments. Because uridine addition resulted in low viability followed by low titre even in its lowest concentration of 2 mM, it was excluded from these set of experiments. Manganese, galactose and FTI were added to concentrations of 25 µM, 40 mM and 15 µM of total culture volume, respectively. Agitation speed and DO set point were set on 190 rpm and 50%, respectively. 2.6% FA and 0.26% FB were added daily from day 2 till day 14 of the batch. Glucose was also supplemented to the culture, daily in order to maintain its concentration about 2.5± 0.2 g/L. All the cultures were stopped on 15^th^ day of the batch.

## Results

### Media and feed selection

Fed batch culture is currently the most common industrial process for CHO cell culture. The aim of fed-batch experiments was to determine the best combination of media/feed that results in the highly optimized viable cell density (VCD), titre and culture longevity (12). Therefore, cell density, viability, final titre and glucose concentration of all studied conditions were compared as represented in Figure 1. The different media and feed combinations that were used in the fed-batch cultures are depicted in Table 1.

**Fig 1.**
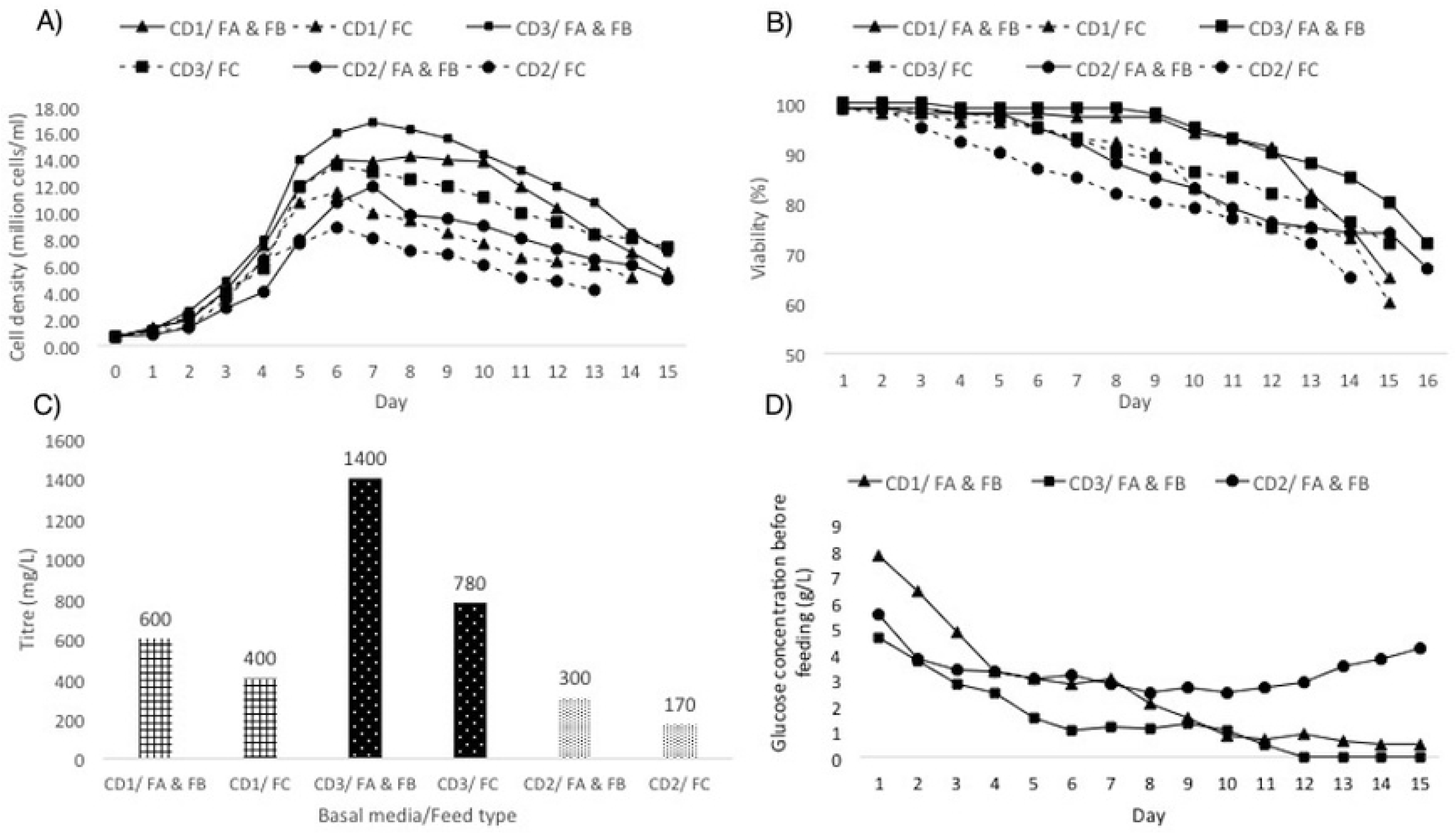
A) Cell density B) Viability, C) Final titre of the different fed-batch experiments performed with different basal/feed combinations and D) Daily Glucose concentration of cultures in different basal media supplemented with FA and FB feeds

CD3 media/FA and FB combination resulted in highest cell density and cell specific productivity of ∽16×10^6^ cells/ml and 8.9 pg/cell/day, respectively. In all three basal media used in this experiment, higher cell and antibody concentrations were obtained when FA and FB feeds were used compared to FC. It has been previously shown that FA and FB are richer than FC both in amino acids and glucose concentration (12). Among basal media used in this experiment, CD2 media resulted in the lowest cell density and final titre irrespective of the feeds that have been used. Otherwise, CD3 media resulted in final titre of 1400 mg/L that was about 2.3 fold higher than CD1, while there was not a significant difference between the cell densities of these two media. Altogether, CD3 media combined with FA and FB feeds were chosen for subsequent experiments on feeding strategy improvement.

As can be seen in Figure 1.B in both CD1 and CD3 media fed with FA and FB feeds, the viability started to drop from day 10 of the batch. Considering daily glucose concentration of these conditions (Figure 1.D), it can be excluded that viability drop may started when glucose level became lower than 1 g/L. This phenomenon also happened when both media were fed with FC feed with the only difference that glucose depletion started earlier (about day 5).

According to the observed results, early viability drop, which coincided with glucose depletion, implies that 1 g/L glucose is a critical limitation concentration for this certain cell line. This result is in agreement with the report suggested using glucose feed in fed-batch process because cells at high density affected by glucose availability (13). In addition, since in these experiments a fixed 2% daily feed was added to the culture, this decreased glucose concentration may reflect the depletion of other nutrients especially amino acids in cultures. Moreover, at time of viability drop the concentrations of both lactate and ammonia were below their inhibition level reported in literature (14).

One of the most efficient feeding strategies is glucose-based feeding. This strategy although can help to maintain glucose levels during the batch but can be problematic due to accumulation or depletion of certain amino acids (12, 15, 16). Meanwhile once-daily feeding strategy cannot ensure that glucose level is being held near the set points pre-defined (13). Based on these results, the decision was made to use fixed feeding strategy while increasing the total amount of daily feed added to the culture. Simultaneously, glucose stock solution of 400 g/L was also used to bring glucose concentration to a level that prevents depletion before the next bolus feed. For this purpose, different glucose set points were also included in the experiments in order to investigate whether this cell line is sensitive to glucose levels or not.

### Integrated medium and feeding strategy optimization

Improvement of cell growth and titre, requires media/feed combinations screening followed by feeding strategy optimization. After choosing CD3 as basal and FA & FB as feeds, RSM design was used to investigate the influence of different feed percentages and glucose set points on productivity of cells. For this purpose, different fixed amount of daily feeds were considered (Table 2). In addition, glucose was fed daily according to the glucose concentration at the moment of feeding and the anticipated glucose uptake rate for the next 24 h. By this method the glucose was held within a pre-determined set point irrespective of the amount of feed added daily. Table 4 shows the final titre of the 13 generated experimental runs after execution.

**Table 4.** Experimental conditions and the titre responses of RSM design.

| Run No. | A: Feed Percent (%) | B: Glc set point (g/L) | Response: Titre (mg/L) |
| --- | --- | --- | --- |
| 1 | 1 | 0 | 2200 |
| 2 | 0 | 0 | 2600 |
| 3 | 1 | -1 | 2400 |
| 4 | 0 | 0 | 2600 |
| 5 | 1 | 0 | 2200 |
| 6 | 0 | 0 | 2380 |
| 7 | -1 | 1 | 800 |
| 8 | 0 | -1 | 2300 |
| 9 | 0 | 1 | 800 |
| 10 | 0 | 0 | 2540 |
| 11 | 1 | 1 | 700 |
| 12 | -1 | -1 | 1600 |
| 13 | 0 | 0 | 2500 |

Factors are represented as their coded values.

The results showed that the max VCC (viable cell count) of 16.5 ± 0.5 ×10^6^ cells/ml was achieved in all experiments. In conditions that Glucose concentration was maintained on 4 g/L, the highest lactate level (50-55 mM), lowest titre (950 ± 250 mg/L) and batch duration (12 days) were observed. It has been previously shown that by increasing the glucose concentration in fed batch mode cultures, higher lactate was produced which indirectly can cause inhibitory effects on growth and productivity (13, 15, 17).

When the glucose concentration of culture was decreased to both 2 and 3 g/L, the lactate consumption was observed after day 4 of the batch and lactate concentration was remained below 10 mM for the rest of the batch. This lower lactate level lead to extended batch duration of 3-days and increased final titre of 2 to 2.5 fold compared to high glucose set point limit of 4 g/L.

In glucose set point of 2 g/L, the final titre increased about 1.4 fold when feed percentage increased from 2% to 2.5 and 3%. It highlights the effect of other nutrients increment in productivity of the culture. In addition, the results also revealed that when cultures are fed using lower amounts of feed (2% daily), increasing the glucose set point level could not significantly affect the final titre. The lower titre seems to be dependent more on other nutrients especially amino acids rather than glucose. When considering the effect of both feeding percentage and glucose set point, the results confirmed that they are in close relationship. The highest final titre of 2.5 ± 0.1 g/L was observed in conditions that were supplemented with 2.5 % daily feed while their glucose set point was adjusted on 3 g/L.

The analysis of variance (ANOVA) results and the proposed model for the final mAb titre response is expressed as an empirical two order polynomial equation in terms of two variables (A: feed percent and B: glucose set point) in Table 5. The larger *F-value* and the smaller *p-value*, show more significant of the corresponding coefficient. The *F-value* of 51.38 indicates that the model is significant. Additionally, in this model feed percent and glucose set point (A and B) linear effects and quadratic effects (A^2^ and B^2^) are significant model terms due to their *P-value* which is lower than 0.05. Besides, interaction between feed percent and glucose set point has also significant effect on protein titre. In addition, the lack of fit *p-value* (0.065), also suggests that the model can be used as a prediction tool.

**Table 5.**
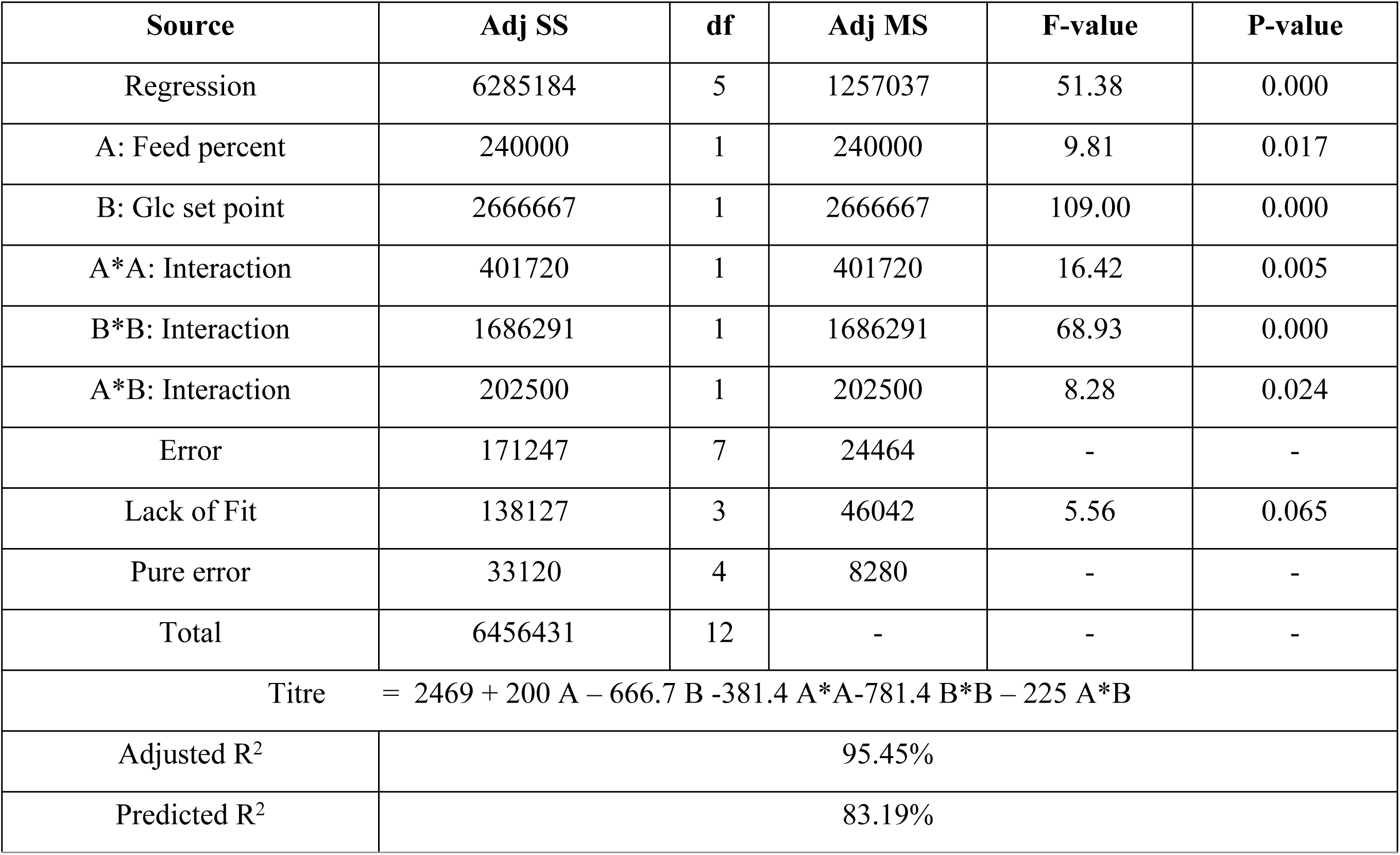
ANOVA results for the Model being built for Titre response in RSM statistical design.

| Source | Adj SS | df | Adj MS | F-value | P-value |
| --- | --- | --- | --- | --- | --- |
| Regression | 6285184 | 5 | 1257037 | 51.38 | 0.000 |
| A: Feed percent | 240000 | 1 | 240000 | 9.81 | 0.017 |
| B: Glc set point | 2666667 | 1 | 2666667 | 109.00 | 0.000 |
| A*A: Interaction | 401720 | 1 | 401720 | 16.42 | 0.005 |
| B*B: Interaction | 1686291 | 1 | 1686291 | 68.93 | 0.000 |
| A*B: Interaction | 202500 | 1 | 202500 | 8.28 | 0.024 |
| Error | 171247 | 7 | 24464 | - | - |
| Lack of Fit | 138127 | 3 | 46042 | 5.56 | 0.065 |
| Pure error | 33120 | 4 | 8280 | - | - |
| Total | 6456431 | 12 | - | - | - |
| Titre = $2469 + 200 A - 666.7 B - 381.4 A*A - 781.4 B*B - 225 A*B$ | | | | | |
| Adjusted R<sup>2</sup> | 95.45% |  |  |  |  |
| Predicted R<sup>2</sup> | 83.19% |  |  |  |  |

The analysis of adjusted and predicted coefficients of determination (R^2^-values) which should be more than 60 indicates that titre response can be easily described with the model being created. (18) Adjusted R^2^ is an indicator of how well the experimental results fit the model while the predicted R^2^ indicates the prediction potential of the model regarding future experiments. Regression analysis shows that the model being built for titre response have adequately high adjusted and predicted R^2^-values (95.45% and 83.19%, respectively).

As can be seen in Table 4, increasing the feed percentage from 2.5% to 3% when glucose set point is on 2 g/L had no significant effect on titre. However in the same scenario when glucose set point was adjusted on 3 g/L, final titre was decreased. Lower titre may be due to the accumulation of specific amino acids which showed inhibitory effect on final productivity.

The results from this experiment shed light in the feeding strategy based on glucose set point for the fed-batch culture of the cell line used in this study. According to the variables coefficients of the model acquired from RSM, main (B) and quadratic (B^2^) effects of glucose set point are the most important parameters affecting the final titre. The other factors can be categorized with order of A^2^ (quadratic effect of feed percent), A (main effect of feed percent) and A*B (interaction effect of feed and glucose set point). This is mainly due to the decreased productivity that has been occurred when increasing the glucose set point to 4 g/L in all conditions irrespective of the amount of feed added. It can be concluded from contour plot (Fig. 2) that regions of increased titre occurred in regions of sufficient amino acids feed (more than 2.5%) with glucose set point of 2.5 ± 0.5 g/L. Senger et al. were also reported that by increasing the glucose set point to values more than 3 g/L especially when the cells were in lack of amino acids, the productivity of r-tPA was decreased (19).

**Fig 2.**
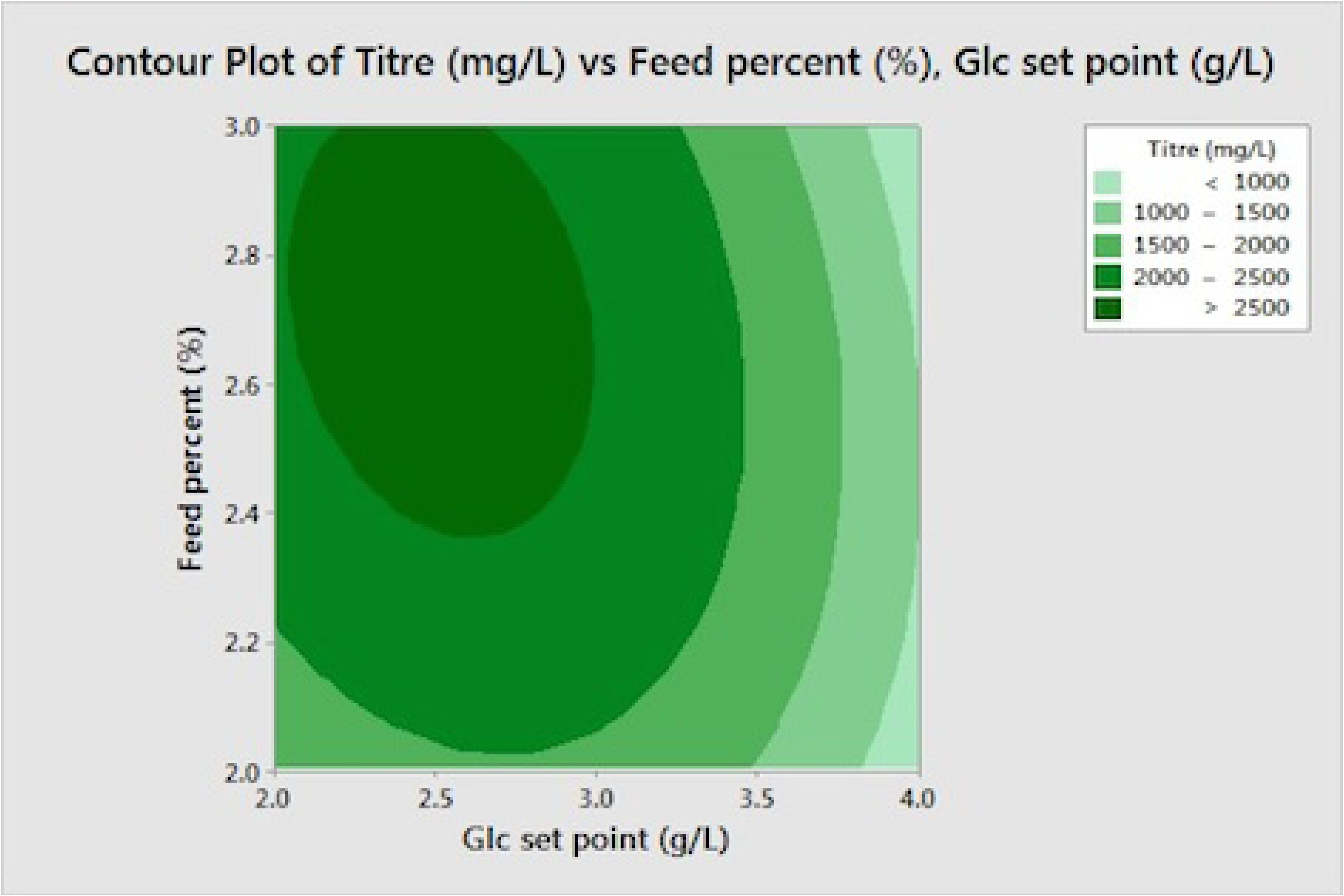
Contour plot for titre of the cultures versus glucose set point and feed percentage.

### Experimental designs regarding glycan variation

In order to improve the glycan structure of trastuzumab biosimilar monoclonal antibody regarding its glycosimilarity with Herceptin originator drug, a set of experiments were designed for increasing the galactosylated and afucosylated glycan structures. Sum of galactosylated and afucosylated oligosacharides in originator drug is usually between 32-35% and 5-9% of total glycan structures.

Galactosylation of Fc region in monoclonal antibodies can affect their complement dependent cytotoxicity (CDC) machanism of action (20, 21). Galactosylation can drop over the course of culture due to reduced intracellular biosynthesis. Inside the cells Uridine triphosphate (UTP) and galactose-1-phosphate react to each other to produce UDP-galactose (UDP-gal) that is the building block of the galactose in oligosacharide chains. The enzyme galactosyltransferase add UDP-gal to the sugar chain and Manganese (Mn^2+^) is the cofactor of this enzyme which helps it to improve its performance. Supplementation of Uridine, manganese and galactose can lead to increased level of UDP-gal that can cause to an increase in galactosylated glycan structures of monoclonal antibodies (11, 22).

Another important Fc-mediated immune effector function that plays the most important role in depleting tumor cells is called antibody-dependent cellular cytotoxicity (ADCC). It has been proved previously that the absence of core fucose on Fc N-glycan structures can lead to enhanced ADCC activity (23, 24). It has also been confirmed that apart from afucosylation, galactosylation levels could also influence ADCC activity; however, the role of afucosylation is more prominent. (25)

Different glycan engineering strategies were used for the purpose of improving the performance of therapeutic monoclonal antibodies through their effector functions (both ADCC and CDC) (26). Supplementation of the media with one of the components involved in galactosylation of monoclonal antibodies namely, uridine (U), mangannese (M) and galactose (G) have provided remarkable results ragarding increased protein oligosacharide galactose content (27-29). Other researchers also demonstrated that adding a mixture of galactose, manganese chloride (MnCl_2_), and uridine to cell culture medium can alter the glycosylation of monoclonal antibodies (11, 22, 30-32).

Different technologies were also applied for reduction of core fucosylation in therapeutic monoclonal antibodies. These include: use of cell lines with inherent reduced capacity for incorporation of fucose (33) and generation of completely non-fucosylated antibodies in cells that were genetically enginered in their *FUT8* gene encoding the α1,6-fucosyltransferase (34). The most straight forward approach that can easily be applied in biopharmaceutical companies is supplementation of media with Fucosyltransferase inhibitors such as 2F-peracetyl-fucose (10).

Given the possibilities discussed above, for improving glycosylation of antibody, combined effect of four factors that can be expected to potentially enhance simultaneously both galactosylation and afucosylation, namely U, M, G and FTI were considered. High and low limits were set for all these factors according to literature and our preliminary studies regarding this antibody. Full factorial design of experiments was used in order to find the most effective factors and also their optimum concentration. This method of DOE was used since it was important to study the impact of several factors on a response simultaneously (35).

The low and high limit of uridine concentration was set on 0 and 4 mM. These concentrations were chosen based on literature reports and subsequently preliminary experiments which showed that cell viability and final titre were remarkably droped when 8 mM uridine was added to culture (data was not shown). Although 4 mM uridine was resulted in lower titre compared to the conditions without this supplement, but in order to check the interaction effect (mainly synergystic effect) of this supplement with manganese and galactose this concentration was considered as the high level in our experiments. Gramer et al. reported that galactose was started to accumulate in culture when it was more than 40 mM (in condition with 8X concentration of UMG). They also confirmed cultures supplemented with 8 mM uridine, 16 µM Mn^2+^ and 40 mM galactose reached to a plateau in their galactosylation level (11). Based on their report and according to our prior knowledge, galactose higher limit was set to 50 mM in our experimental design. The high limit of Mn^2+^ concentration was chosen according to the patent No. US20170107551A1 in which they reported that 40 µM manganese concentration resulted in the most optimum galactosylated level in shake flasks (36). David T.Ho et al. results represented that using 100 µM FTI resulted in 67% decrease in afucosylation level (10). Besides, in preliminary experiments with FTI supplement, it was observed that adding 50 µM FTI in culture resulted in a decreased fucosylation level from 97 ±1% (in absence of FTI) to 85 ±1 %. According to this experiment, the decision was made to consider 0 and 30 µM FTI concentrations as the lowest and highest level for FTI factor for finding its omptimum concentration.

When galactose was added to culture with concentration of 50 mM it could increase the galactosylated glycans about 8% in absence of both Mn^2+^ and Uridine supplements (Fig 3). About 11% increase in galactosylation level was also reported when 20 mM galactose was added to CHO fed-batch cultures (37).

**Fig 3.**
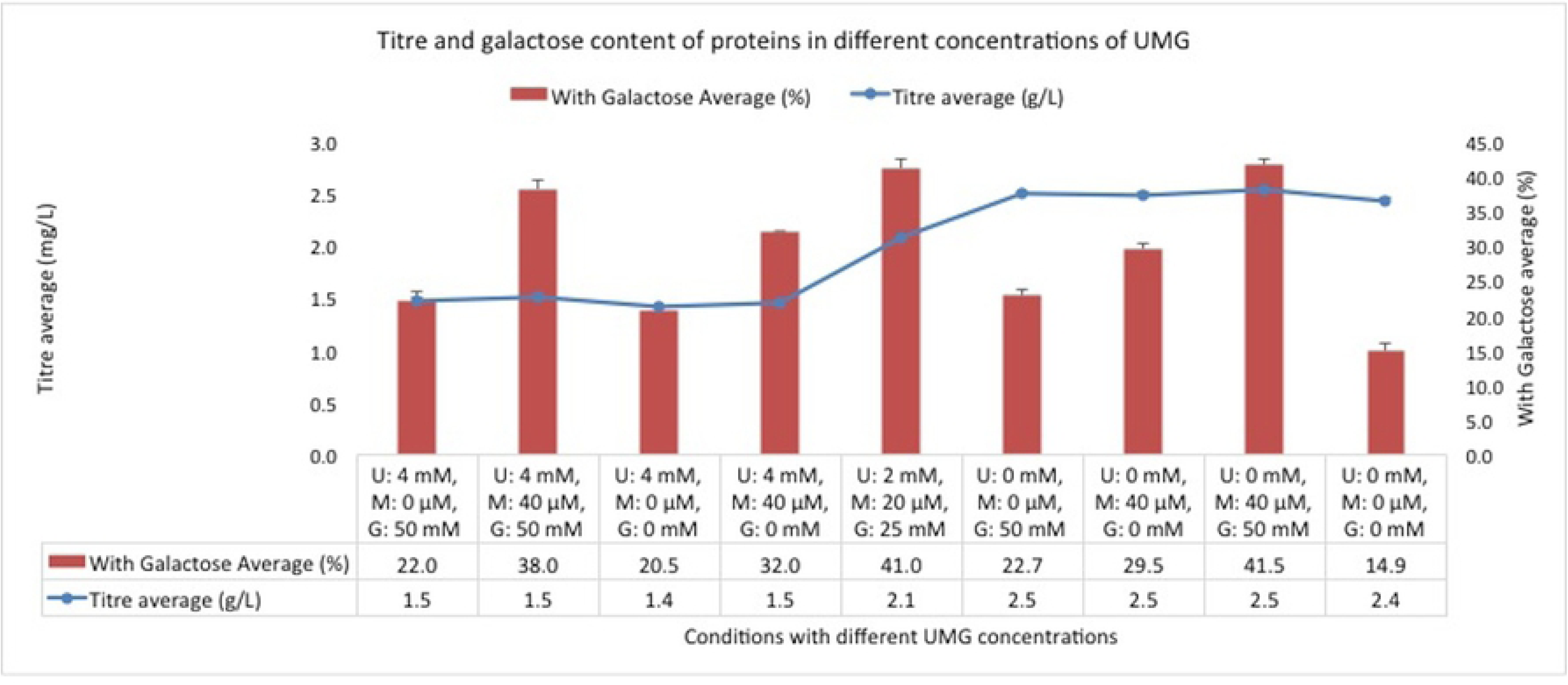
Titre and galactosylation level of different experimental runs with different concentrations of UMG. The data are represented as mean ± SD.

Addition of 4 mM uridine to the culture containing 50 mM galactose couldn’t further increase the level of galactosylation. Furtheremore in presence of 4 mM uridine lower cell density was observed followed by lower final titre of 1.5 ±0.2 g/L compared to the titre of about 2.5 g/L in absence of uridine. It was in contrast to Gramer et al. reports in which they showed that addition of uridine up to 16 mM had no effect on productivity reduction (11). Our results indicated that addition of Mn^2+^ led to increased protein oligosacharide galactose content from 22±0.5% (in absence of Mn^2+^ and presence of galactose) to 30±2% (in absence of galactose) and to 40±2% (in presence of either 25 or 50 mM galactose).

The overall increase in galactosylation was primarily due to a drop in G0F content with a related increase in G1F and G1F’ and secondarily to a slight increase in G2F (Table 7). As can be seen in Table 5 the G0F content of the monoclonal antibody was about 70% in absence of Mn^2+^ and decreased to 48% and 58% in presence of 20 and 40 µM Mn^2+^, respectively. G1F glycan structures was increased from 9% (in absence of Mn^2+^ and galactose) to 14.5 ± 0.5% (in absence of Mn^2+^ and presence of galactose) and to 25 ± 1% (in presence of both Mn^2+^ and galactose). The trend of increase in G1F’ structures was exactly similar to the G1F glycans with the only difference that the increase in these oligosacharides were happened with lower slope. This is in total agreement with the results expained by Roger Anderson et al., in which they claimed that galactosylation of glycan structures preferentially occures more on the glycans α1,6 than α1,3 arm (38). G2F glycans were only increased about 3 ±0.5% when both Mn^2+^ and galactose were supplemented in cultures while there was no change in these glycans when either Mn^2+^ or galactose were present.

**Table 6.**
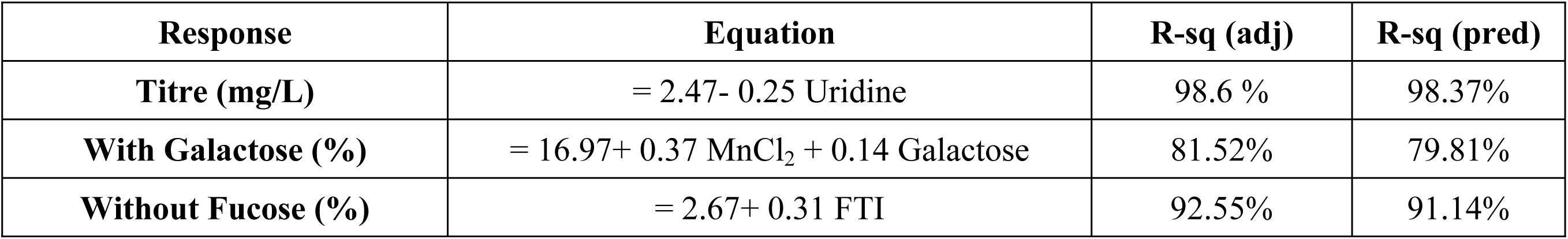
Linear regression models obtained from factorial design statistical analysis.

**Table 7.** Detail of glycan structure percentage in different concentrations of uridine, manganese and galactose in absence and presence of FTI.

| U (mM), M (μM) and G (mM) | FTI (μM) | G0 | G0F | G1 | G1' | G1F | G1F' | G2F | Galactosylation (%) | Afucosylation (%) |
| --- | --- | --- | --- | --- | --- | --- | --- | --- | --- | --- |
| **4, 0, 50** | 0 | 3.3 | 67.2 | 0.55 | 0.4 | 14.7 | 4.3 | 3 | 23 | 4.3 |
|  | 30 | 10.6 | 64.4 | 0.9 | 0.5 | 13 | 3.8 | 2.8 | 21 | 12 |
| **4, 40, 50** | 0 | 1.9 | 55.5 | 0.8 | 0.7 | 25.3 | 7.7 | 4.5 | 39 | 3.4 |
|  | 30 | 11.6 | 47.4 | 0.86 | 0.5 | 22.9 | 8.3 | 4.4 | 37 | 13 |
| **4, 0, 0** | 0 | 1.6 | 72.9 | 0.5 | 0.4 | 13.1 | 3 | 2 | 19 | 2.5 |
|  | 30 | 12.6 | 65.3 | 1.7 | 0.7 | 11.5 | 4.1 | 2 | 20 | 15 |
| **4, 40, 0** | 0 | 2.7 | 61.9 | 0.2 | 0.03 | 23.6 | 5 | 1.2 | 30 | 3 |
|  | 30 | 8.7 | 53.6 | 0.86 | 0.44 | 18 | 8.3 | 4.37 | 32 | 10 |
| **2, 20, 25** | **15** | **8.1** | **48.7** | **1.3** | **0.04** | **26** | **8** | **4.6** | **40** | **9.5** |
| **0, 0, 50** | 0 | 1.1 | 69 | 0.5 | 0.4 | 15.2 | 4.3 | 3 | 23 | 2 |
|  | 30 | 10.6 | 63.4 | 0.9 | 0.5 | 13.8 | 4 | 2.8 | 22 | 12 |
| **0, 40, 0** | 0 | 1.2 | 63.4 | 0.2 | 0.03 | 23.6 | 5 | 1.2 | 30 | 1.5 |
|  | 30 | 9.8 | 57 | 0.8 | 0.4 | 18.5 | 5.8 | 3.5 | 29 | 11 |
| **0, 40, 50** | 0 | 1.2 | 52.4 | 0.46 | 0.02 | 26 | 9.5 | 5.96 | 42 | 1.7 |
|  | 30 | 9.7 | 47 | 0.86 | 0.44 | 24.1 | 8.5 | 5 | 39 | 11 |
| **0, 0, 0** | 0 | 1.7 | 77.3 | 0.18 | 0.06 | 9 | 3.3 | 1.41 | 14 | 2 |
|  | 30 | 10.9 | 68.5 | 1.54 | 0.04 | 8 | 3.9 | 1.5 | 15 | 12.5 |

G0F-GN and Man5 structures do not included in the table since their percentage was less than 4% and 3%, respectively in different experiments.

The equations of the models and the main effect plots (derived in Minitab) for all three responses are shown in Table 6 and Figure 4, respectively. These models were fitted by omitting unsignificant terms based on their *p-values* (>0.05). All three responses have linear relationships with their effective factors. As represented in Table 6, the models gained from factorial design analysis also revealed that the percentage of galactosylated glycans was mostly dependent on MnCl_2_ followed by galactose concentrations with coefficients of 0.37 and 0.14, respectively. The results obtained were completely in agreement with Crowell et al. study in which they represented that addition of Mn^2+^ alone can lead to increased protein oligosacharide content (29). The negative coefficient of uridine concentration in titre response also revealed that its presence in culture negatively affect the productivity of the cell line used in this study.

**Fig 4.**
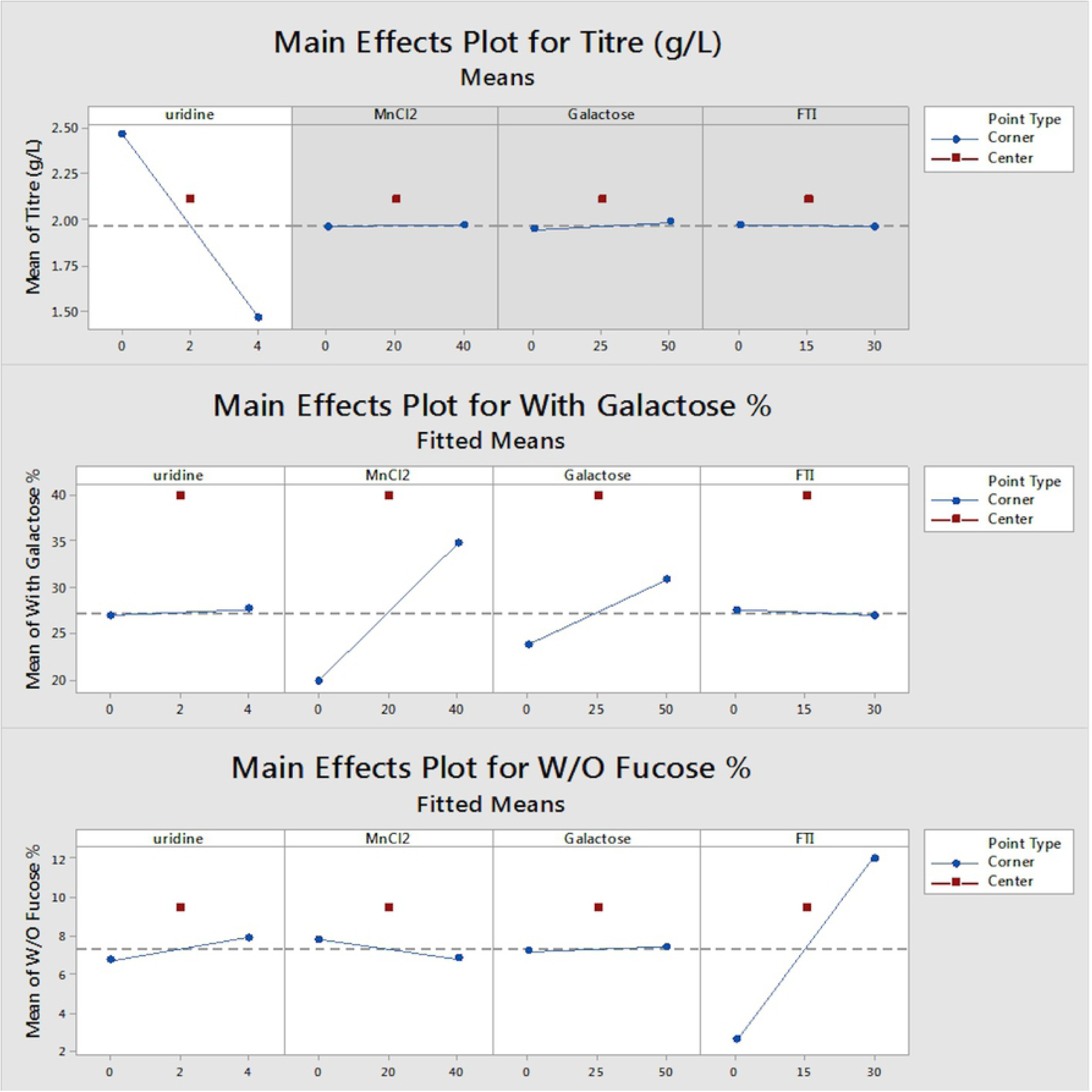
Main effects plot of titre, with galactose % and W/O Fucose% responses Vs, uridine, MnCl_2,_ galactose and FTI factors in factorial design analysis.

In the context of afucosylation, our results confirmed that using FTI as a fucosyltransferase inhibitor we could increase the afucosylated glycan structures of trastuzumab biosimilar about 8%. This increase in afucosylated structures was similar in both FTI concentrations (15 and 30 µM) that were used in our experiments. Interestingly, it can also be concluded from the results that the increase in afucosylated structures happened irrespective of the presence of other three supplements used for galactosylation improvement. Furtheremore, supplementation of the inhibitor to the culture had not influenced culture parameters in terms of viable cell density and productivity, exactly the same as David T.Ho results (10). Figure 4 also confirmed that FTI supplement is the only factor that is responsible for increasing the afucosylated glycans level. The main increase in afucosylated glycan content was due to the increase in G0 content of oligosacharides being present in trastuzumab biosimilar drug which was increased from about 2 ± 1% in conditions without FTI to 8 ± 1% and 9.5 ± 1% in presence of 15 and 30 µM FTI, respectively.

It also should be noted that afucosylation was slightly increased from 1.8 ±0.2% (in absence of uridine) to 3.5 ± 1% (in presence of 4 mM uridine). Nevertheless, in presence of uridine even in its lowest amount (2 mM), lower titre was observed compared to absence of this supplement. Gramer et al. also reported that the fucosylation of their antibody encountered with a slight drop from 97% to 94% in 0 and 20 mM uridine (11). The main effect plot (Fig. 4A) also shows that the most significant of the investigated parameters which affects titre was uridine concentration. In addition negative coefficient of uridine concentration in titre response equation indicates that it negatively affects titre. Due to this negative impact on culture behaviour and according to its negligible effect on afucosylation glycan level, the decision was made not to supplement this factor in future cultures. The supplementation of U, M and G supplements neither affected Man 5 nor fucosylated glycan content. Besides, other quality attributes of the protein have not been affected by supplementing the fed-batch cultures with all these four supplements.

Our results pointed out that control on the amount of both galactosylation and afucosylation content of monoclonal antibodies being produced in CHO cell lines can be achieved by supplementing the culture simultaneously with UMG and FTI. Taken together, these experiments suggests that the highly optimized concentration of Mn^2+^, galactose and FTI for simultaneously controlling both the galactosylation and afucosylation level of glycan structures were 20 µM and 25 mM, and 15 µM, respectively. Furthermore, supplementing the cultures with all these glycosylation modulators neither affect the culture performance and productivity nor other quality attributes of the protein regarding its comparability with originator drug.

### Verification and validation of models

For demonstrating the predictive ability of both RSM and factorial designs obtained respectively for feeding strategy and glycan improvement, a set of experiments were performed with the optimum level of the factors acquired from response optimizer of Minitab software. The feeding strategy and supplementation patterns used in verification experiments are represented in Table 8. The verification process generated experimental results close to the predicted response (*p-value* < 0.05) as represented in Table 9. Our optimized feeding strategy combined with glucose, galactose, MnCl_2_ and FTI supplementation resulted in mean expression of 2510 ± 65 mg/L. Simultaneously the level of galactosylation and afucosylation were also increased to 32.5 ± 0.6 % and 7.3 ± 0.5 %, respectively which were completely comparable to Herceptin originator drug. The good agreement between the experimental and predicted results verifies the validity of the models.

**Table 8.** Optimal processing conditions used for model verification in 5-Lscale bioreactors.

|  |  |
| --- | --- |
| **Feeding strategy** | 2.6 % daily FA+ 0.26% FB daily. Start feed: day 2 |
| **Glucose set point (g/L)** | Glucose was supplemented to culture in a way to maintain glucose above $2.5 \pm 0.2$ g/L |
| **Galactose (mM)** | 40 |
| **$\text{MnCl}_2$ ($\mu\text{M}$)** | 25 |
| **FTI ($\mu\text{M}$)** | 15 |

**Table 9.** Comparison of the predicted versus experimental results obtained in model verification experiments.

|  | **Titre (mg/L)** | **With galactose %** | **Without Fucose %** |
| --- | --- | --- | --- |
| Experimental value of 5 L scale runs | $2510 \pm 65.19$ | $32.5 \pm 0.62$ | $7.37 \pm 0.57$ |
| Predicted value | 2600 | 32 | 7.44 |
| Significant? (Y/N) | N* | N | N |

N: not significant

## Conclusion

The focus of this study is not only on improving the process titre but also on product quality and comparability. The aim of this study regarding process yield was to investigate the effects of different basal media and feeds on cell culture performance and mAb production. The highest mAb titre (1.4 g/L) was obtained with CD3 basal medium and FA and FB feeds. In next phase of development, the feeding strategy was optimized through high throughput response surface methodology which resulted in a fixed feeding strategy on a daily basis while controlling the glucose level of culture (more than 2 g/L on the next day of culture). By operating with a feed and glucose control, higher titre of about 2.5 ± 0.1 g/L were achieved. The feeding strategy that was developed was beneficial to our company since it was easy to implement this strategy both in lab-scale and in manufacturing. Another goal of this study regarding product quality was achieved through supplementation of culture with glycan structures precursors or glycosyltransferase enzymes regulators. Although the glycan structures regulation can be achieved using commercial available supplements such as EX-CELL Glycosylation Adjust which enables the desired N-linked glycosylation, combination of supplements have been chosen to be investigated that can not only modulate glycan structures but also can be cost beneficial when used as a cocktail in large scale production. To achieve this, full factorial design of experiments were included to find the effect of UMG and FTI supplements on titre, galactosylation, and afucosylation content of trastuzumab biosimilar drug. Among supplements used, the best combination achieved was 25 µM Mn^2+^, 40 mM galactose and 15 µM FTI. These supplements resulted in galactosylation level of more than 30% and afucosylation of 5-9% which is completely similar to its originator. To our knowledge, this study is the first report regarding the application of both afucosylation and galactosylation improving supplements for glycan engineering of pharmaceutical proteins. This work provided a methodology that can be used for media and process development studies. Future studies might involve adding these supplements at different time points during culture to determine and specify the best time for their supplementation.

## Abbreviations

mAb: Monoclonal antibody;
DOE: design of experiment;
RSM: response surface methodology;
CDC: complement dependent cytotoxicity;
ADCC: antibody dependent cell cytotoxicity;
CQA: critical quality attributes;
Fc: fragment crystallizable;
FTI: fucosyl transferase inhibitor;
CHO: Chinese hamster ovary;
HER2: human epidermal growth factor receptor 2;
CCD: central composite design;
ANOVA: analysis of variance;
HPLC: high pressure liquid chromatography;
STR: stirred tank reactor;
DO: Dissolved oxygen;
VCD: viable cell density;
VCC: viable cell count.

## Acknowledgements

This work was supported by Biopharmaceutical Research Center of AryoGen pharmed Inc. and was performed under ethical supervision of Alborz University of Medical Sciences with ethics code of IR.ABZUMS.REC.1396.042 and IR.ABZUMS.REC.1397.036.

